# Patterns of thaumarchaeal gene expression in culture and diverse marine environments

**DOI:** 10.1101/175141

**Authors:** Paul Carini, Christopher L. Dupont, Alyson E. Santoro

## Abstract

Thaumarchaea are ubiquitous in marine habitats where they participate in carbon and nitrogen cycling. Although metatranscriptomes suggest thaumarchaea are active microbes in marine waters, we understand little about how thaumarchaeal gene expression patterns relate to substrate utilization and activity. Here, we report the global transcriptional response of the marine ammonia-oxidizing thaumarchaeon ‘*Candidatus* Nitrosopelagicus brevis’ str. CN25 to ammonia limitation using RNA-Seq. We further describe the genome and transcriptome of *Ca*. N. brevis str. U25, a new strain capable of urea utilization. Ammonia limitation in CN25 resulted in reduced expression of transcripts coding for ammonia oxidation proteins, and increased expression of a gene coding an Hsp20-like chaperone. Despite significantly different transcript abundances across treatments, two ammonia monooxygenase subunits (*amoAB*), a nitrite reductase (*nirK*), and both ammonium transporter genes were always among the most abundant transcripts, regardless of growth state. *Ca*. N. brevis str. U25 cells expressed a urea transporter 139-fold more than the urease catalytic subunit *ureC*. Gene co-expression networks derived from culture transcriptomes and ten thaumarchaea-enriched metatranscriptomes revealed a high degree of correlated gene expression across disparate environmental conditions and identified a module of genes, including *amoABC* and *nirK*, that we hypothesize to represent the core ammonia oxidation machinery.

**Originality-Significance Statement:** Discovering gene function in fastidious or uncultivated lineages remains one of the biggest challenges in environmental microbiology. Here, we use an approach that combines controlled laboratory experiments with *in situ* transcript abundance data from the environment to identify genes that share similar transcription patterns in marine ammonia-oxidizing thaumarchaea. These findings demonstrate how transcriptomes from microbial cultures can be used together with complex environmental samples to identify suites of co-expressed genes that are otherwise enigmatic and provide new insights into the mechanism of ammonia oxidation. Our results add to the growing body of literature showing that relatively small changes in transcript abundance are linked to large changes in growth in organisms with reduced genomes, suggesting they have limited capacity for metabolic regulation or that they rely on mechanisms other than transcriptional regulation to deal with a fluctuating environment.

## Introduction

Ammonia-oxidizing thaumarchaea are ubiquitous and abundant in the oceans, accounting for >30% of all cells below the thermocline (Karner *et al*., 2001; Schattenhofer *et al*., 2009) and are integral organisms in oxygen minimum zones (Francis *et al*., 2005; Coolen *et al*., 2007; Lam *et al*., 2009; Pitcher *et al*., 2011; Stewart *et al*., 2012; Beman *et al*., 2012). In many marine environments, thaumarchaeal transcripts are among the most abundant that can be mapped to available prokaryotic genomes (Hollibaugh *et al*., 2011; Baker *et al*., 2012; Stewart *et al*., 2012; Gifford *et al*., 2013). In these environments, the most frequently detected thaumarchaeal transcripts encode for proteins involved in ammonia oxidation and acquisition, including ammonia monooxygenase subunits (*amoABC*), ammonium transporters (*amtB*), a putative Cu-containing nitrite reductase (*nirK*), and structural cellular components (for example, S-layer proteins (Nakagawa and Stahl, 2013)). In addition to dissolved ammonia, some ammonia-oxidizing archaea utilize ammonia derived from urease-catalyzed urea hydrolysis as a chemolithoautotrophic growth substrate (Qin *et al*., 2014; Bayer *et al*., 2015) and urease genes and transcripts believed to be of thaumarchaeal origin have been detected in marine environments (Shi *et al*., 2010; Alonso-Sáez *et al*., 2012; Pedneault *et al*., 2014; Tolar, Ross, *et al*., 2016). Despite the abundance of thaumarchaeal transcripts in natural assemblages, we still have a poor understanding of how the relative abundance of thaumarchaeal transcript markers such as *amoA, nirK* and *ureC* relate to nutrient and energy availability across environmental gradients.

Thaumarchaeal ammonia oxidation is initiated by the oxidation of ammonia to hydroxylamine (NH_2_OH) by the ammonia monooxygenase enzyme complex (Amo) (Vajrala *et al*., 2013), but the enzyme(s) catalyzing the oxidation of NH_2_OH to nitrite (NO_2_ ^−^) have not been confirmed (Walker *et al*., 2010). Orthologs of the bacterial hydroxylamine oxidoreductase (Hao) or c-type cytochrome synthesis and assembly machinery, thought to be required for NH_2_OH oxidation and electron transfer in ammonia-oxidizing bacteria (AOB) (Arp *et al*., 2007), are absent from all sequenced thaumarchaeal genomes (Stahl and la Torre, 2012; Spang *et al*., 2012; Kerou *et al*., 2016). Instead, unidentified Cu-containing metalloenzymes or F_420_-dependent monooxygenases are speculated to be involved in NH_2_OH oxidation and electron transfer to archaeal terminal oxidases (Walker *et al*., 2010; Kerou *et al*., 2016) and may involve nitric oxide (NO) as either a direct intermediate or an electron shuttle (Stieglmeier *et al*., 2014; Martens-Habbena *et al*., 2015; Kozlowski *et al*., 2016). While the precursor to NO has not yet been elucidated, all free-living thaumarchaea with complete genomes encode a Cu-containing multicopper oxidase with homology to Cu-dependent nitrite reductases (NirK) that may be responsible for the reduction of NO_2_ ^−^ to NO (Kerou *et al*., 2016).

‘*Candidatus* Nitrosopelagicus brevis’ str. CN25 is a cultured representative of ubiquitous and abundant pelagic thaumarchaeal populations in the shallow oligotrophic ocean (Santoro and Casciotti, 2011; Santoro *et al*., 2015). Here, we describe the genome and transcriptome during urea-based growth of a *Ca*. N. brevis strain that can utilize ammonia cleaved from urea as a sole chemolithoautotrophic growth substrate. Additionally, we use *Ca*. N. brevis str. CN25 to investigate the transcriptional response to ammonia limitation in laboratory culture. These transcriptomes are further analyzed in the context of several marine metatranscriptomes and used to identify conserved gene co-expression networks.

## Results and Discussion

### ‘*Candidatus* Nitrosopelagicus brevis’ strain U25 genome and transcriptome

A urea-utilizing thaumarchaeon was obtained from an ammonia-oxidizing enrichment culture (Santoro and Casciotti, 2011) via subculturing with urea as a sole nitrogen and energy source (see ‘Experimental Procedures’) and used for the shotgun metagenome sequencing and experiments described here. After assembly and contig binning based on nucleotide frequencies and coverage, we obtained a three contig genome of this urea-utilizing thaumarchaeon (Supplementary Figure 1a). This genome is nearly identical to the *Ca*. N. brevis CN25 genome with regards to *i)* gene content; *ii)* genome organization (Supplementary Figure 1b); and *iii)* genome wide average nucleotide identity (99.99%; Supplementary Figure 2). We found 19 additional genes at four distinct genomic loci in this strain, relative to CN25 (Supplementary Table 1). The largest of these insertions (15 contiguous genes) includes 11 genes coding urea utilization machinery, including *ureABCDEFG*, which codes for urease and its chaperones, two urea sodium:solute symporter family (SSSF) transporters, a transcriptional regulator and several hypothetical proteins (Fig. 1a). We designate this urea-utilizing thaumarchaeon ‘*Candidatus* Nitrosopelagicus brevis’ strain U25.

**Figure 1:**
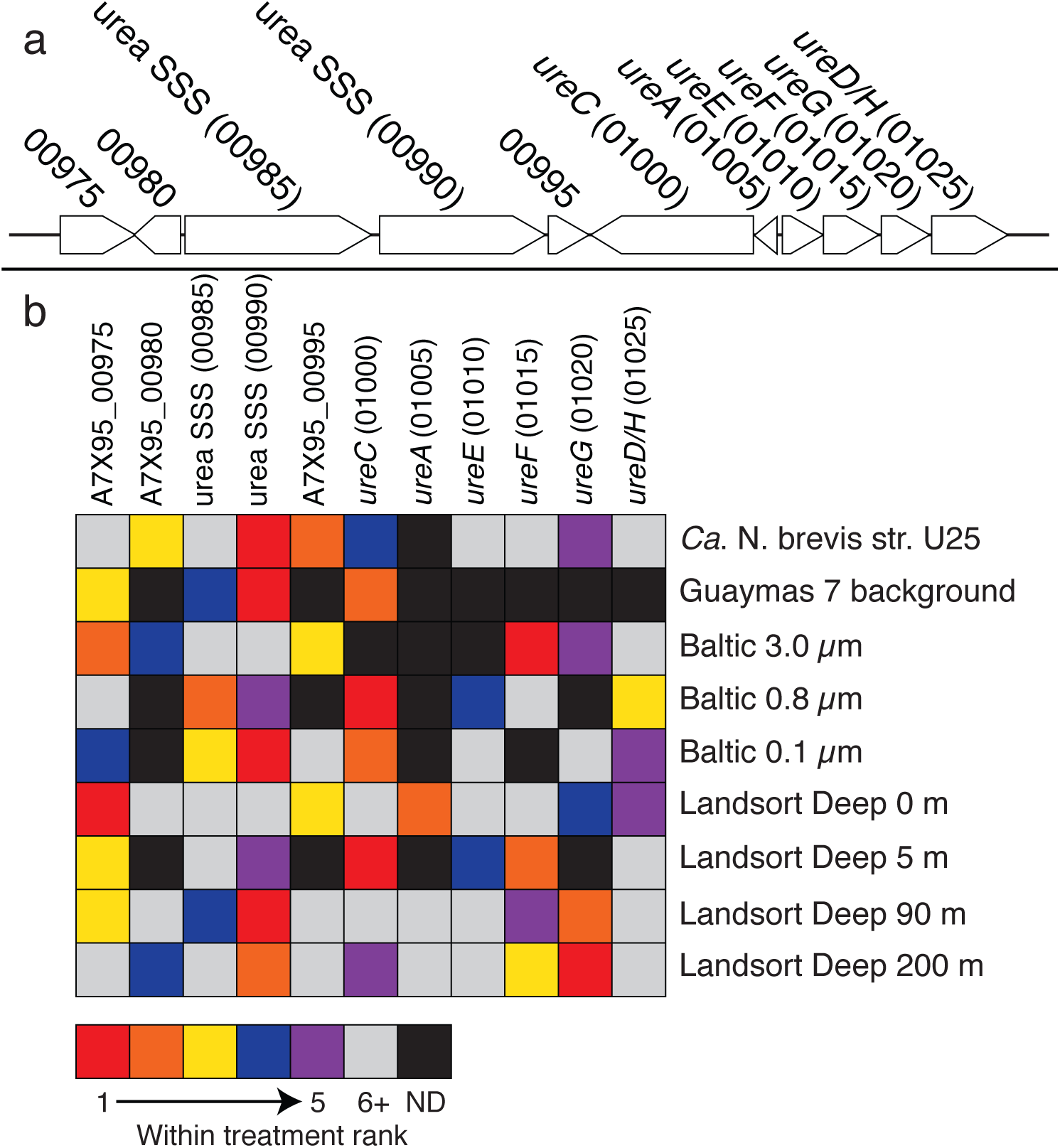
Urea transport and catalysis genes are frequently the most highly expressed *Ca*. N. brevis U25-specific genes. (a) Chromosomal orientation of the *Ca*. N. brevis str. U25 indel conferring urea transport and catalysis capability. (b) Heatmap illustrating the relative rank expression level of the five most abundant genes within the urea indel region for culture experiments and environmental metatranscriptomes. Rank was calculated within a given treatment from RPKM normalized expression values, where a rank of 1 is the most abundant transcript of the genes contained in the indel. The mean expression (n=3 replicates) value was used for ranking the culture treatment. ND=Not detected. Note this is not the rank of the transcript within the entire metatranscriptome. Some metatranscriptomes were excluded because they did not have sufficient coverage of the urea transport and catabolism machinery. Numbers in parentheses after gene names refer to A7X95 locus tags.

**Figure 2:**
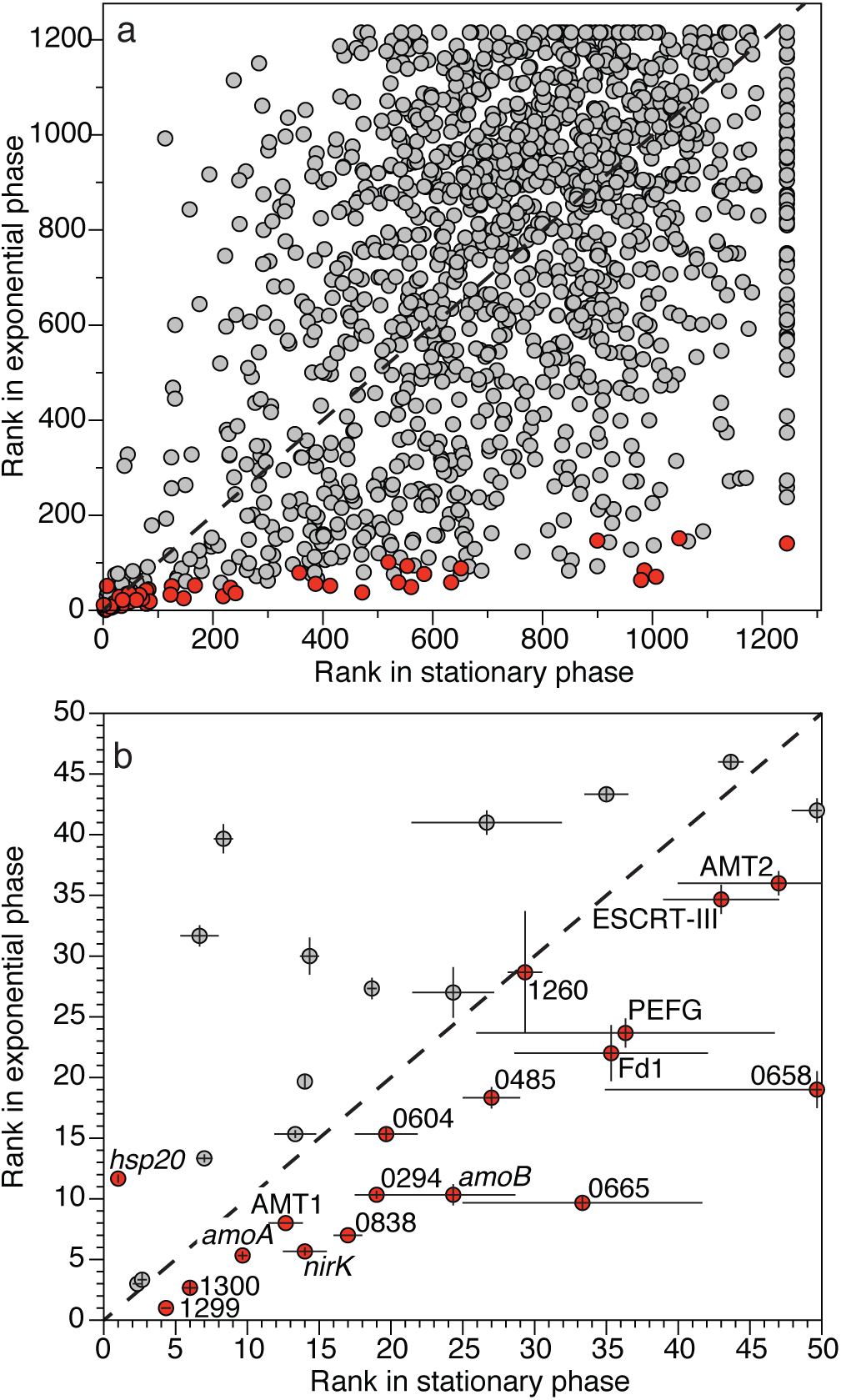
Highly expressed *Ca*. N. brevis str. CN25 transcripts in exponential phase are also highly expressed in stationary phase, despite significant differences in abundance. (a) Points are the mean rank (n=3) of RPKM normalized expression values for all genes in exponential and stationary growth phases. Red points are transcripts that were significantly differentially abundant across treatments (Supplementary Table 2). The abundances of grey points were not significantly different across treatments. Dashed line is 1:1 line indicating no change in rank. (b) Subset of panel (a), illustrating the rank of transcripts that are in the top 50 most abundant transcripts in both exponential and stationary phase. Points in (b) are the mean rank and error bars represent ± SE (n=3). Gene transcript abundances in (b) that were significantly different across treatments are labeled and colored red. The ‘T478_’ prefix is omitted from labels of genes annotated as ‘hypothetical’. PEFG corresponds to T478_0596.

We sequenced a transcriptome from *Ca*. N. brevis str. U25 growing exponentially with urea as the growth substrate. Only one transcript from the chromosomal insertion containing the urea transport and metabolism genes (Fig. 1a) was among the top 50 transcripts detected: A7X95_00990, coding for a putative urea SSSF transporter (ranked 13.7 ± 0.33; mean ± SE, *n* = 3). Surprisingly, transcripts coding for catalytic urease components, or the second putative urea SSSF transporter (A7X95_00985) located immediately adjacent to A7X95_00990, were not nearly as abundant as A7X95_00990. For example, the mean expression levels of *ureC*, coding for the fused catalytic αβ-subunit of urease, and *ureA* (γ urease subunit) were 139 and 784-fold less abundant than that of A7X95_00990, respectively (ranked 390 ± 19.7 and 950 ± 50.8, respectively; mean ± SD, *n*=3). A7X95_00985, coding for the second urea SSSF transporter, was also expressed at a low level, comparable to *ureA*, ranked 940 ± 92.0. During growth on urea, transcripts for genes coding for an ammonium transporter (AMT1), ammonia monooxygenase subunits (*amoAB*) and nitrite reductase (*nirK*) were within the top ten most abundant transcripts detected in strain U25.

Although urea utilization genes have been detected in wild thaumarchaeal populations, we have a poor understanding of how the abundances of urease transcripts relate to growth and activity. To contextualize the expression patterns observed in U25, we compared the relative rank of the transcript abundances for only those genes coding for urea uptake and catabolism (Fig. 1a) under laboratory growth conditions to the relative rank of the transcript abundances of the same genes from several deeply sequenced marine metatranscriptomes. The SSSF urea transporter (A7X95_00990) was the most abundant urea-related gene transcript in 38% of the environmental datasets (*n* = 8) we investigated (Fig. 1b). In contrast to culture conditions, where the SSSF urea transporter was the most abundant urea-related gene transcript, *ureC* was the most abundant transcript in 25% of the environmental datasets (Fig. 1b). This shows that variability in the relative transcriptional activity of urea transport and catabolism genes is not unusual. Our finding that *ureC* was not highly expressed in exponentially growing cells also helps to explain previous field observations of low *ureC* expression, and suggests the abundance of *ureC* transcripts may be a poor molecular biomarker of active urea-based nitrification. For example, *ureC* expression and urea-based nitrification were found to be only weakly correlated across several environments (Tolar, Wallsgrove, *et al*., 2016), in contrast to high correlation between *amoA* expression and ammonia oxidation rates (J. M. Smith *et al*., 2014). Similarly, in Arctic samples collected across seasons, *ureC* genes were detected, yet *ureC* transcripts were only sporadically detected and at low abundances (Pedneault *et al*., 2014).

### Transcriptional response to ammonia limitation in ‘*Ca*. N. brevis’ strain CN25

To understand adaptive mechanisms during ammonia starvation, we explored the transcriptional response of *Ca*. N. brevis CN25 to ammonia limitation. A total of 51 gene transcripts were differentially abundant when comparing the exponential growth phase of CN25 to ammonia-limited stationary phase (generalized linear model likelihood ratio test FDR ≤ 0.01 and ≥ 2-fold change in abundance; Supplementary Table 2). The gene transcripts that were significantly less abundant in stationary phase included *amoA, amoB, nirK*, both *amtB*-like ammonium transporters (AMT1=T478_1378; AMT2=T478_1350), several additional Cu-containing metalloenzymes, and two ferredoxin-like 4Fe-4S binding domain proteins (Fd1=T478_1472 and Fd2=T478_1259) (Fig. 2, Supplementary Table 2). The downregulation of *amoA, amoB* and *nirK* in ammonia-limiting conditions was recently shown for the ammonia-oxidizing thaumarchaeon *Nitrosopumilus maritimus* using DNA microarrays (Qin *et al*., 2017), suggesting that one adaptation of ammonia-oxidizing archaea to ammonia limitation is to reduce the relative expression of energy generation machinery.

Only two genes, T478_1481, coding an Hsp20/α-crystallin domain small heat shock protein, and T478_1394, annotated as a hypothetical protein, were significantly more abundant (∼10-fold) in ammonia-limited stationary phase (Fig. 2, Supplementary Table 2). Hsp20 is a molecular chaperone that enhances thermotolerance and binds to unfolded proteins to prevent aggregation (Li *et al*., 2011). The higher proportion of Hsp20 transcripts and concomitant decrease in proportion of transcripts coding enzymes integral to energy production suggests that one adaptation *Ca*. N. brevis employs in ammonia-limited stationary phase may be to protect existing proteins from degradation. Nutrient stress has been shown to induce the expression of molecular chaperone proteins in ammonia-oxidizing archaea, ammonia-oxidizing bacteria, and oligotrophic marine heterotrophs. For example, in *N. maritimus*, two copies of Hsp20 were differentially expressed during copper stress and recovery, but not during ammonia starvation (Qin *et al*., 2017). Similar to our findings regarding Hsp20, *Nitrosomonas europaea* expressed peptides for the molecular chaperone GroEL in both energy starved and energy replete conditions, but at significantly greater levels under energy starvation (Pellitteri-Hahn *et al*., 2011). The authors speculated that energy stress may induce chaperone expression as part of a generalized stress response, and that these chaperones are involved in protein protection (Pellitteri-Hahn *et al*., 2011). Similarly, the marine chemoorganoheterotroph ‘*Candidatus* Pelagibacter ubique’ induced GroEL protein expression under N-starvation (D. P. Smith *et al*., 2013), GroES under iron starvation (D. P. Smith *et al*., 2010), and the heat shock protein IpbA in nutrient-limited stationary phase (D. P. Smith *et al*., 2016). These finding suggest that one role of molecular chaperones during nutrient stress may be to protect key enzymes from proteolytic turnover when cells scavenge peptides to support nutrient-limited sustenance.

In contrast to the finding that only two genes were more abundant in ammonia-limited stationary phase for *Ca*. N. brevis, over 200 genes were upregulated in ammonia-limited stationary phase for *N. maritimus* (Qin *et al*., 2017). One explanation for this observation is that *Ca*. N. brevis has a reduced capacity to sense and respond to environmental change as a result of its small genome (Santoro *et al*., 2015). Alternatively, our conservative statistical thresholding may have resulted in a reduced number of genes defined as differentially expressed. While it is important to note that our experimental design cannot distinguish between responses to energy limitation versus anabolic nitrogen limitation, transcriptional responses to ammonium limitation that involved the upregulation of only a few genes has been described for both ammonia-oxidizing bacteria and oligotrophic marine bacteria. For example, in the bacterium *N. europaea*, only 0.42% of the genome was upregulated under N-starvation (Wei *et al*., 2006). Similarly, ‘*Ca*. P. ubique,’ exhibited a weak transcriptional response to N-starvation (D. P. Smith *et al*., 2013). The similar lack of transcriptional responses to nitrogen starvation by *Ca*. N. brevis and *Ca*. P. ubique are consistent with observations that organisms with small and potentially streamlined genomes have a limited capacity to respond rapidly to environmental change (Giovannoni *et al*., 2014; Cottrell and Kirchman, 2016; Giovannoni, 2017; Satinsky *et al*., 2017).

Despite significantly different transcript abundances of essential ammonia oxidation and transport genes across growth conditions, the rank order of these transcripts within a given treatment were similar, irrespective of growth condition. In particular, the most abundant gene transcripts in exponential phase were generally still the most abundant transcripts in ammonium-limited stationary phase (Fig. 2b). For example, transcripts for 32 genes (64%) were in the top 50 most abundant transcripts in both exponential and stationary phase (Fig. 2b). Interestingly, the abundances of 18 of these transcripts were also significantly different across treatments (Fig. 2b), illustrating that although transcripts can be differentially abundant across paired treatments, the changes in their relative cellular abundance may be subtler. Several of these consistently abundant but differentially expressed transcripts are common molecular markers predicted to be essential for ammonia oxidation and transport, including the ammonium transporters AMT1 and AMT2, ammonia monooxygenase subunits (*amoAB*) and nitrite reductase (*nirK*). This suggests that the proportional changes we observed in highly-expressed genes may be the result of abundance changes in other genes that make up a smaller proportion of the transcriptome, such as Hsp20, and that even though we observe significant differences across treatments, the underlying transcript abundances might be similar.

The ammonia monooxygenase C subunit (AmoC) has been implicated in stress response (Berube and Stahl, 2012) and recovery from ammonia starvation in *N. europaea* (Berube *et al*., 2007). The participation of AmoC in ammonium starvation appears to be conserved in thaumarchaeal ammonia oxidizers, where *amoC* transcript levels remained high in ammonia-limited stationary phase (Qin *et al*., 2017). Consistent with these findings, *amoC* was abundant in both exponential and N-limited stationary phase in CN25 (the 13^th^ and 7^th^ most abundant transcript, respectively), and we did not observe a significant difference in the abundance of *amoC* across growth phases (Fig. 2).

### Correlated gene expression across disparate environments

Controlled laboratory experiments such as those described above help us to understand gene regulation by isolating one experimental variable at a time. However, gene expression patterns observed in natural thaumarchaeal populations are the result of cells responding to complex and dynamic environmental conditions that can be cryptic and difficult to mimic in the laboratory. To identify clusters of co-expressed genes across disparate environmental conditions, and relate them to our laboratory findings, we constructed and analyzed a gene expression correlation network constructed from transcriptomes of exponentially growing CN25 and U25 cultures and marine metatranscriptomes. Although 64 metatranscriptomes were mapped to the *Ca*. N. brevis genomes, only ten met our strict criteria for inclusion in the network analysis presented here (see ‘Experimental Procedures’, below). The transcription of 1,407 of the 1,464 non-redundant genes in the two *Ca*. N. brevis genomes was significantly positively correlated with at least one other gene (Pearson’s r ≥ 0.80, *q* ≤ 0.025; Fig. 3, Supplementary Table 3). Network modularity is a measure of the group connectivity within a network, where connections contained within a module are denser than connections between modules. Modularity values range from −0.5 to 1, where 1 describes a highly modular system. The modularity of this positive correlation network was 0.71, indicating a high degree of modularity. Genes with positively correlated expression organized into 38 groups (modules), ranging in membership size from 2 to 236 genes (mean module size = 38.0 genes).

**Figure 3:**
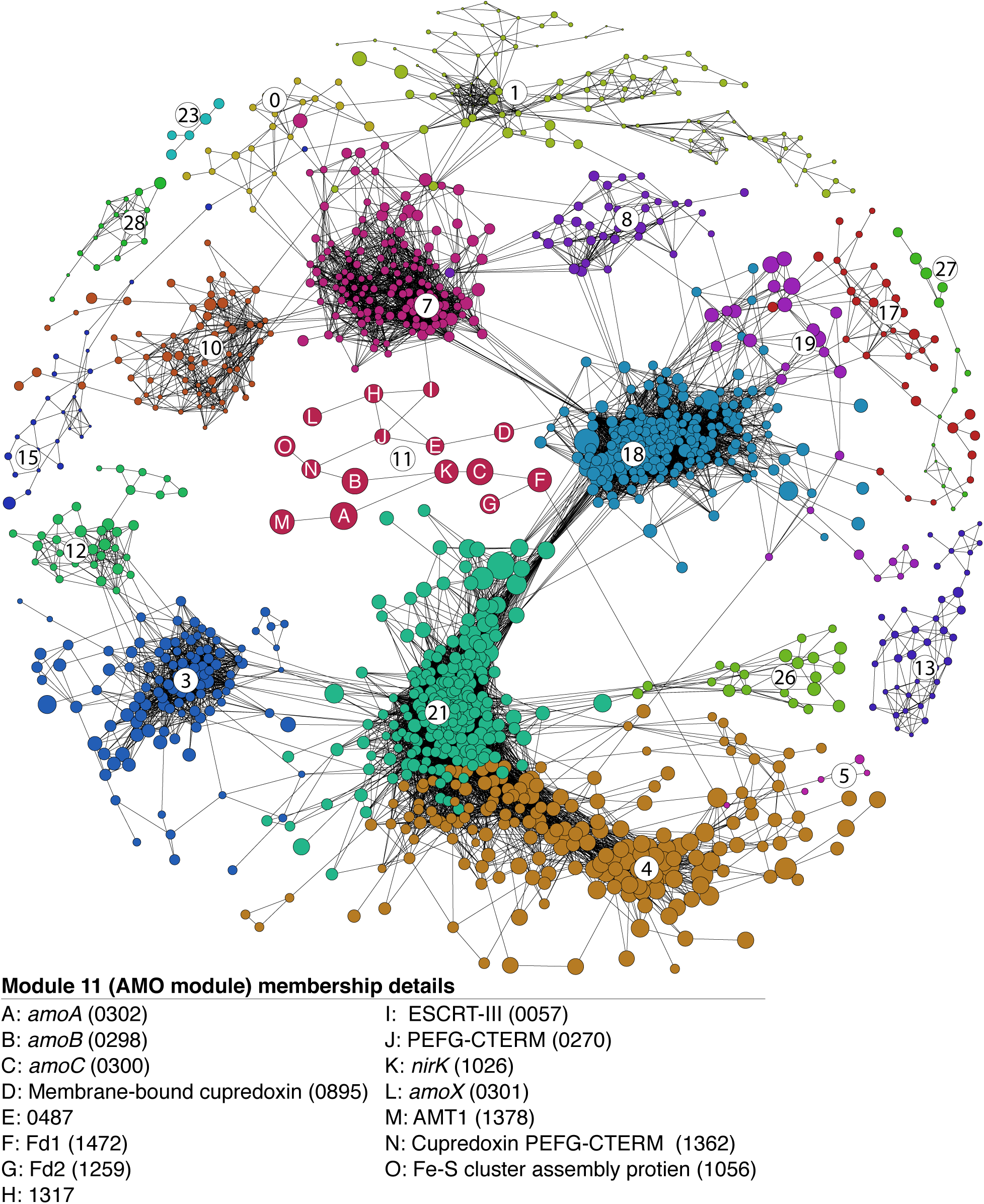
Thaumarchaeal gene expression is highly modular and ammonia oxidation genes are co-expressed. Network diagram of strong and significant (Pearson’s r ≥ 0.8, *q* value ≤ 0.025) positive correlations across ten environmental metatranscriptomes and the *Ca*. N. brevis U25 transcriptome depicted in Fig. 1 and a *Ca*. N. brevis CN25 transcriptome from a culture initiated with 100 µM NH_4_Cl (12 conditions total, see methods). Individual nodes are genes. Nodes are sized by the mean normalized rank abundance (VdW scores), whereby larger nodes are more abundant transcripts on average. Nodes are colored by module membership. Circled numbers are the module identity (Supplementary Table 3). The module 11 (the AMO module) genes are identified by letters A-O; their annotations are provided below network. Numbers in parentheses refer to T478 locus tags. Only modules with five or more nodes are shown for clarity; see Supplementary Table 3 for all module membership details.

Genes encoding putative components of the core ammonia oxidation and transport machinery are significantly co-expressed across distinct environmental and laboratory conditions. A single 15-gene module (module 11 in Fig. 3) contained: *amoABCX*, AMT1, *nirK*, two PEFG-CTERM domain proteins, Fd1 and Fd2, an Fe-S cluster assembly protein, a membrane bound cupredoxin-containing protein, and three hypothetical proteins. Further investigation of these hypothetical proteins suggests that (T478_0057) is a putative archaeal cell division protein related to the endosomal sorting complexes involved in membrane trafficking (ESCRT)-III (Lindås *et al*., 2008; Spang *et al*., 2015). Our finding that *amoA, amoB* and *nirK* transcripts are abundant and co-expressed with other genes coding for membrane-bound Cu-containing metalloproteins (T478_1362 and T478_0895) implies the products of these genes may participate in ammonia oxidation. Previous speculation implicated membrane-bound multicopper oxidases (Walker *et al*., 2010; Stahl and la Torre, 2012; Kozlowski *et al*., 2016) or novel F_420_-dependent monooxygenases (Kerou *et al*., 2016) in ammonia oxidation chemolithotrophy (specifically NH_2_OH oxidation) based on Cu redox chemistry or ortholog conservation across thaumarchaeal genomes. However, the genes put forth in those studies were not present in module 11 (referred to as the AMO module, herein), suggesting they may not be involved in core energy metabolism (Fig. 3, Supplementary Table 3). For example, a multicopper oxidase present in the genomes of both *N. brevis* (T478_0261) and *N. maritimus* (Nmar_1663) previously implicated to be involved in the oxidation of hydroxylamine to nitrite (Walker *et al*., 2010; Qin *et al*., 2017) was not present in the AMO module (Supplementary Table 3).

On average, the AMO module is expressed at a higher level than other modules, and was more abundant in conditions where ammonia oxidation rates and thaumarchaeal abundances would be predicted to be high (Fig. 4). For example, consistent with previous reports of higher ammonia oxidation rates within hydrothermal plumes (Lam *et al*., 2004), we show that the AMO module is expressed highly within the Guyamas Deep hydrothermal plume, relative to background samples (Fig. 4). Moreover, similar to reports of increased thaumarchaeal gene expression in the mesopelagic (Church *et al*., 2010), the AMO module is less abundant in the surface waters of Landsort Deep (0 and 5 m), relative to deeper waters (90 and 200 m) (Fig. 4).

**Fig. 4:**
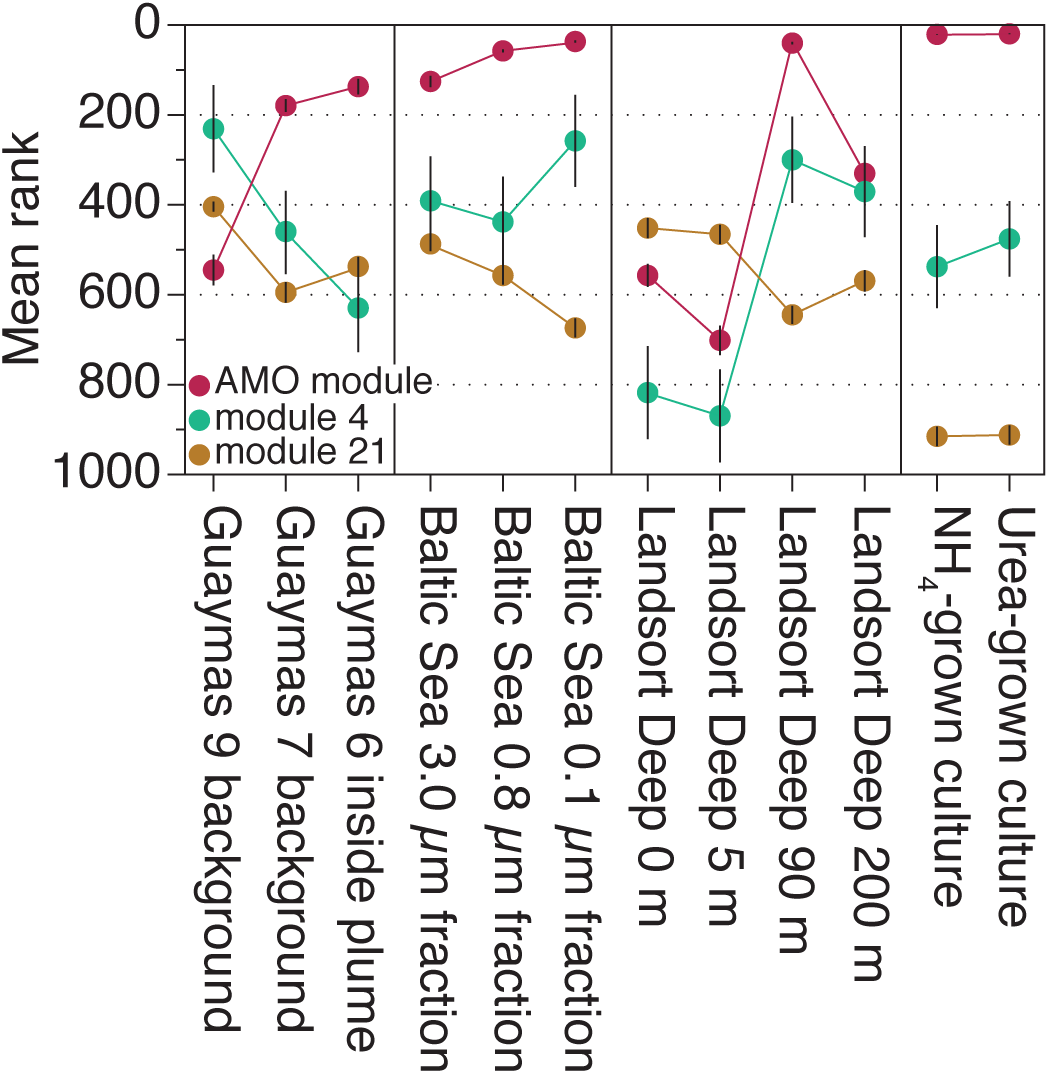
Natural variability in the mean rank of expression modules across environments and culture conditions. Points are the average rank of all genes contained within a given module at each site; error bars are ± SE. On average, the AMO module and modules 36, 4 and 21 rank the highest (that is, they are the most abundant) across all sites. Module 36 is not displayed because it is only comprised of two genes, which prevents statistical analysis, and these genes are absent in the U25 genome.

### A new proposed model for thaumarchaeal chemolithotrophy via ammonia oxidation

A previous model of thaumarchaeal ammonia oxidation proposed NO is derived from the reduction of NO_2_^−^ by NirK, and that this NO is subsequently used to oxidize NH_2_OH (Kozlowski *et al*., 2016). However, this model does not agree with tracer experiments using ^18^O labeled water, which show that only one O atom from water is incorporated into NO_2_^−^ produced by thaumarchaea (Santoro *et al*., 2011; Buchwald *et al*., 2012). If NO_2_^−^ was reduced to NO and used to produce additional NO_2_^−^, the resulting NO_2_^−^ would retain an average of more than one O atom from water. Based on the co-expression data presented here, we propose two alternative models of thaumarchaeal ammonia oxidation that are consistent with previous isotope tracer data regarding the source of O atoms in NO_2_^−^. In both models, NirK and two membrane-anchored cupredoxins (T478_1362 and T478_0895) act in concert to oxidize NH_2_OH to NO_2_^−^ in two steps: a three-electron oxidation of NH_2_OH to NO, followed by a one-electron oxidation of NO to NO_2_^−^ (Supplementary Figure 3). The involvement of T478_0895 orthologs in electron transfer with NirK, is consistent with previous predictions based on gene expression data for *N. maritimus* (Qin *et al*., 2017). Both proposed pathways are also consistent with the observed co-expression and high abundance of these transcripts across distinct environmental conditions (Figs 2-4; (Hollibaugh *et al*., 2011; Baker *et al*., 2012; Stewart *et al*., 2012; Gifford *et al*., 2013)). The predicted localization of ammonia oxidation in the pseudoperiplasm (Walker *et al*., 2010) is also a key aspect of the proposed models, as protein domain analysis with InterPro (Jones *et al*., 2014) indicates all proteins in these models, and pertinent cupredoxin domains, are likely localized in the pseudoperiplasmic space. While we cannot determine which reaction is conducted by which Cu metalloenzyme, both scenarios are more parsimonious than existing models and are plausible based on existing bioinorganic chemistry literature.

Similar to the *N. maritimus* transcriptome and environmental metatranscriptomes (Hollibaugh *et al*., 2011; Williams *et al*., 2012; Qin *et al*., 2017), ferredoxins were among the most highly expressed genes in *N. brevis* (Fig. 2). However, we were surprised to find these ferredoxins (Fd1 and Fd2) and an Fe-S cluster assembly protein co-expressed with ammonia oxidation genes in the AMO module. The co-expression of ferredoxins in the AMO module suggests a central role for Fe-S cluster-containing proteins in the electron transport chain of thaumarchaea. Both Fd1 and Fd2 lack discernable signal sequences or PEFG domains, suggesting that they are localized in the cytoplasm. Ferredoxin-containing DNA binding transcriptional regulators have been implicated as NO sensors (Kiley and Beinert, 2003). However, there are no predicted DNA-binding domains in Fd1 or Fd2. Sequence structure threading of Fd1 and Fd2 with phyre2 (Kelley *et al*., 2015) returned best structural matches to NADH dehydrogenase (ubiquinone) iron-sulfur protein 8 (threading confidence score = 99.8% for both Fd1 and Fd2; coverage of Fd1 was 99% and 71% for Fd2). Thus, we speculate that Fd1 and Fd2 participate in supplying electrons to the ubiquinone pool, and thus may be involved in supplying the electrons necessary to initiate ammonia oxidation via Amo.

Microbial gene expression is inherently modular. For example, functionally related genes are often co-expressed in operons or regulons as a response to external environmental signals (Freyre-Gonzales *et al*., 2010). However, the organization and patterns of population-level gene expression in dynamic environments are poorly understood. Although we show a high degree of modularity and correlated expression of a set of genes likely involved in ammonia oxidation, the gene membership of the remaining modules did not reveal a clear pattern of how the genes contained within a given module are functionally related (Supplementary Table 3). For example, genes coding for the modified 3-hydroxypropioinate/ 4-hydroxybutryrate carbon fixation pathway proteins are dispersed across five modules (module numbers 3, 4, 18, 21, and 36; Fig. 3 and Supplementary Table 3). Similarly, vitamin B_12_-production genes are spread across eight modules (module numbers 1, 3, 4, 10, 12, 18, 21, and 26; Fig. 3 and Supplementary Table 3). The structure of the transcription co-expression network illustrated in Fig. 3 is consistent with the interpretation that thaumarchaeal population-level regulatory organization is structured in a decentralized manner. One explanation for this organization may be that decentralized expression networks may help to buffer gene expression changes or maintain genetic diversity in dynamic environments through a suite of feedbacks without centralized regulatory mechanisms (Hartwell *et al*., 1999).

Other reasons the gene membership in the co-expression modules may not reveal clear functional relationships may stem from the environments sampled or the analysis techniques used. For example, our methods likely underestimate the true modularity of *Ca*. N. brevis gene expression. Some of the larger expression modules may comprise distinct modules that we could not resolve because we did not sample an environment with physicochemical parameters necessary to resolve subtle gene expression patterns. Second, although our goal was to be conservative in our network construction, we may be missing important network structural components because of the thresholding parameters or our analysis techniques. Further resolution of such gene expression patterns would require deeply sequenced metatranscriptomes from additional distinct environments and transcriptome analysis of additional thaumarchaea grown under diverse culture conditions. Future research into understanding why certain genes or pathways are co-expressed with one another, and how transcript abundances manifest into functional potential in each environment or culture setting would be necessary to fully disentangle the gene expression we observe here.

## Conclusions

Here we show the rank of most thaumarchaeal transcripts reported as being abundant in the environment (*amoABC*, both *amtB* genes and *nirK*, for example) are relatively invariant across growth phases and environmental conditions. That is, within a given treatment, abundant genes are consistently proportionally abundant, irrespective of growth condition. However, consistent with other studies of thaumarchaeal transcription, the proportions of some of these genes were indeed significantly differentially abundant across paired treatments, indicating ammonia availability did affect the proportional abundances of ammonia oxidation and transport transcripts. One explanation for this observation is that although these transcripts are not ‘constitutive’ in a classic sense (that is, they are differentially abundant across paired experiments), they are instead ‘affluent,’ in that they make up a large part of the total transcript pool, irrespective of growth condition.

Discovering gene function in fastidious or uncultivated lineages remains one of the biggest challenges in environmental microbiology. Narrowing the scope of targets for detailed biochemical investigation is difficult because manipulative experiments are limited in their ability to identify networks of co-regulated genes by the number of environmental parameters we can recreate in a laboratory. The approach used here leverages *in situ* transcript abundance data - in which the environmental conditions are incompletely characterized - to identify genes that share similar transcription patterns. In addition to our putative models of ammonia oxidation in thaumarchaea, this approach shows that 4Fe-4S cluster-containing proteins likely have an important role in ammonia oxidation, indicating a role for iron in archaeal nitrification, which has been previously under appreciated. Detailed biochemical characterization of NirK, other cupredoxin-containing proteins, Fd1 and Fd2 is the next step in understanding their specific role in core thaumarchaeal energy metabolism.

## Methods

### Organism sources

Both ‘*Candidatus* Nitrosopelagicus brevis’ str. CN25 and ‘*Ca*. N. brevis’ str. U25 were obtained from an ammonia-oxidizing enrichment culture previously known as CN25 (Santoro and Casciotti, 2011) grown on Oligotrophic North Pacific Medium. Oligotrophic North Pacific Medium (ONP) consists of natural seawater, a chemolithoautotrophic nitrogen source (NH_4_Cl or urea), ampicillin (10.8 μM), streptomycin (68.6 μM), potassium phosphate (29.4 μM), and a chelated trace metal mix consisting of disodium ethylenediaminetetraacetic acid (14 μM), FeCl_2_ (7.25 μM), ZnCl_2_ (0.5 μM), MnCl_2_ (0.5 μM), H_3_BO_3_ (1 μM), CoCl_2_-6H_2_O (0.8 μM), CuCl_2_-2H_2_O (0.1 μM), NiCl_2_-H_2_O (0.1 μM), Na_2_MoO_4_-2H_2_O (0.15 μM). Preliminary metagenomic sequencing of the original CN25 enrichment indicated the presence of urease genes in a minority of the archaeal population. Sequential transfers of the initial enrichment were made into ONP (Santoro and Casciotti, 2011) amended with 50-100 μM urea, instead of NH_4_Cl, over a period of ∼48 months. In parallel, separate transfers of the CN25 enrichment culture were propagated using NH_4_Cl as a nitrogen and energy source, but without the use of streptomycin, which could potentially serve as a source of urea (Klein and Pramer, 1961). The enrichment culture resulting from the propagation with NH_4_Cl did not contain amplifiable *ureC* genes by PCR, using the archaeal-specific ureC primers Thaum_UreC F and Thaum_UreC R (Yakimov *et al*., 2011) before the sequencing and genome analysis described elsewhere (Santoro *et al*., 2015).

### General cultivation conditions

All thaumarchaeal enrichments were propagated in in ONP medium in a base of aged natural seawater (collected from 10 m depth at 15°S, 173°W on 23 October 2011; 0.2 μm pore size filtered at sea) amended with 50 or 100 μM NH_4_Cl or 100 μM urea as the chemolithoautotrophic substrate. All cultures were propagated in 250 mL polycarbonate flasks at 22°C in the dark and monitored for NO_2_ ^−^ production using the Griess reagent colorimetric method (Strickland and Parsons, 1972). Cell counts were obtained with a Millipore Guava EasyCyte 5HT flow cytometer as described previously (Tripp, 2008).

### Cell harvesting for and genome sequencing of ‘Ca. N. brevis’ str. U25

The *Ca*. N. brevis U25 enrichment culture that was grown exclusively with urea as the sole chemolithoautotrophic growth substrate for >50 generations, was harvested by filtration on to 25 mm diameter, 0.22 μm pore-size Supor-200 filters and frozen at −80°C. DNA was extracted using a DNeasy blood & Tissue DNA extraction kit (Qiagen, Valencia, CA, USA), following the manufacturer’s instructions. The DNA was treated with RNAse and examined using a Bioanlayzer 2100 (Agilent) with 500 ng serving as the input for library construction (NEBNext paired-end DNA Library Prep kit, New England Biolabs). The sample was sequenced on an Illumina MiSEQ (v2 chemistry, paired 250 bp reads). Reads were quality trimmed and served as the inputs to assembly with metaSPAdes (v 0.5, 70mer) (Nurk *et al*., 2017). The K-mer usage and phylogenetic annotation of the assembled contigs were then used to visually identify a putative thaumarchaeal bin (Supplementary Figure 1a) (Laczny *et al*., 2015). The 3 contig genome was annotated using the JGI IMG pipeline (img.jgi.doe.gov) and the PGAP pipeline at NCBI (Zhao *et al*., 2011).

### Experimental design, cell harvesting and RNA extraction for culture transcriptomes

For experiments investigating the effects of ammonia limitation, ONP medium was amended with 50 μM NH_4_Cl as the chemolithoautotrophic growth substrate. In this experiment, six replicates were prepared, three of which were harvested in late exponential phase (Exponential phase) and three of which were harvested in late NH_4_Cl-limited stationary phase (Stationary phase) (Supplementary Fig. 4). We deliberately harvested the exponential phase cultures in late exponential phase to ensure maximal cell biomass for transcriptome analysis. The length of starvation in stationary-phase was based on the exponential-phase doubling time of seven days, approximately the population doubling time during exponential growth. NH_4_Cl-limitation in this phase is supported by a linear dose response in maximal cell density to NH_4_Cl additions (Supplementary Fig. 5).

Transcriptomes that were included in the network analysis were obtained from distinct, mid-exponential phase *Ca*. N. brevis strains CN25 and U25, that were growing on ONP medium amended with either 100 μM NH_4_Cl (str. CN25; n=3) or 100 μM urea (str. U25; n=3) as growth substrates, respectively (Supplementary Fig. 6). Cells were harvested by filtration on to 25 mm diameter, 0.22 μm pore-size Supor-200 filters and frozen at −80°C. For RNA extraction, cells were disrupted as described in (Santoro *et al*., 2010). RNA was extracted using TRIzol LS reagent (Ambion-Life Technologies) per the manufacturer’s instructions and stored in nuclease-free water at −80°C. Urea consumption by str. U25 was determined colorimetrically using the diacetyl monoxime method (Price and Harrison, 1987) (Supplementary Fig. 7).

### Transcriptome sequencing and mapping for culture experiments

Transcriptome samples were prepared for sequencing using the TotalScript RNA-Seq kit (Epicentre-Illumina), which biases against rRNA, using the manufacturer recommended protocol. Libraries were trial sequenced on an Illumina MiSEQ to determine uniformity between barcodes and then fully sequenced in one 300 cycle NextSEQ run which generated 246.6 million paired-end 150 bp reads. Raw Illumina reads in fastq format were interleaved to match paired ends. Sequencing primers and barcode indexes were identified by BLAST against the NCBI vector database and trimmed along with regions with Q scores < 30. Reads mapping to ribosomal RNAs were identified and removed using ribopicker (Schmieder *et al*., 2011). Reads were then mapped to *Ca*. N. brevis genomes at 90% nucleotide identity with CLC Genomics Workbench (command: clc_ref_assemble –s 0.9). Raw read counts per open reading frame (ORF) were compiled.

### Analysis of differentially abundant gene transcripts

Differential gene abundance analysis was performed using a generalized linear model likelihood ratio test in the edgeR software package (v 3.8.5) (Robinson and Smyth, 2008). We defined significant differential abundance as those genes with a false discovery rate (FDR) ≤ 0.01 and greater than 2-fold abundance change across treatments.

### Rank Analyses

Raw read counts per ORF were scaled to expression units of reads per base per million reads mapped (RPKM=(10^6^ * C)/(NL/10^3^)) where C is the number of transcript reads mapped to an ORF; N is total reads mapped to all ORFs in the genome; and L is the ORF length in base pairs (Mortazavi *et al*., 2008). RPKM values were subsequently ranked, with a rank of 1 depicting the most abundant transcript within a given treatment. Rank ties within a treatment were averaged.

### Metatranscriptome mapping to genomes of Ca. N. brevis strains

Sequence reads from 68 metatranscriptomes were mapped to the *Ca*. N. brevis genomes at 50% nucleotide identity using CLC Genomics Workbench (command: clc_ref_assemble –s 0.5) (Supplementary Table 4). The number of metatranscriptome reads that mapped to the *Ca*. N. brevis genomes were variable and ranged from 10 reads to 236,954 reads and mapped to 0.5-89% of the unique genes in the *Ca*. N. brevis genomes (Supplementary Table 4). Raw read counts per ORF were then compiled (ORF n=1445 for str. CN25 and n=1461 for U25).

### Network construction

Only those metatranscriptomes that mapped to ≥45% of the ORFs in the *Ca*. N. brevis genomes, along with the transcriptomes from *Ca*. N. brevis strains CN25 and U25 growing in exponential phase initiated with 100 μM NH_4_Cl or urea, respectively, were included for network analysis. Of the 68 metatranscriptomes mapped to the *Ca*. N. brevis genomes, only ten passed this filtering step and were used for network analysis (Supplementary Table 4). Read counts were scaled to RPKM expression units. RPKM scores calculated for individual culture transcriptome replicates were averaged (n=3) to avoid pseudo-replication effects in the network. The resulting RPKM expression values were rank-normalized to Van der Waerden (VdW) scores using the formula (*s* = Φ^−1^(*r* / (*n*+1))), where *s* is the VdW score for a gene, *r* is the rank for that observation, *n* is the sample size and Φ is the Φ^th^ quantile from the standard normal distribution using *tRank* in the multic R package (Lunde *et al*.). Pearson correlation coefficients and *P* value estimates were calculated for all gene:gene pairs across the VdW-normalized metatranscriptomes and culture experiments (n=10 and 2, respectively) with the *rcorr* command in the Hmisc R package (Harrell and Dupont). To correct for multiple hypothesis testing, *q* values were computed from *P* value estimates using the qvalue R package (Storey and Tibshirani, 2003). Correlations with a *q* value ≤ 0.025 were used for network analysis. All correlations at this threshold were strongly correlated (Pearson’s *r* ≥ 0.8).

### Network Statistics

Network modularity and module membership were calculated in Gephi (0.8.2 beta) with the following settings: resolution 1.0, randomized and unweighted (Blondel *et al*., 2008). The resulting network was visualized using the Fruchterman-Reingold algorithm in Gephi.

## Data Availability/Sources

Transcriptomes from *Ca*. N. brevis str. CN25 and U25 can be found in the NCBI BioSample archive under accession numbers SAMN6290440-6290457. The U25 genome has been deposited at DDBJ/ENA/GenBank under the accession LXWN00000000. The version described in this paper is version LXWN01000000. The metatranscriptomic data from Landsort Deep in the Baltic is available from the Sequence Read Archive under numbers SAMN04943349-SAMN04943415. Other metatranscriptomes analyzed in the network are publically available through iMicrobe (https://imicrobe.us) or NCBI’s Short Read Archive through the following accession numbers: CAM_PROJ_Sapelo2008, CAM_PROJ_AmazonRiverPlume, CAM_PROJ_PacificOcean, CAM_P_0000545, SRA023632.1.

## Acknowledgements

This research was supported by NSF awards OCE-1260006, OCE-1437310, and DBI-1318455 to AES. CLD was supported by NSF OCE-1259994. We thank Matt Rawls for urea measurements, Mike Stukel for obtaining oligotrophic seawater, and Albert Barberán and Jason Corwin for discussions regarding network construction. AES is an Alfred P. Sloan Research Fellow in Ocean Sciences.

