## Supplementary Materials for "Patterns of thaumarchaeal gene expression in culture and diverse marine environments"

### Supplemental Information for:
### Correlated expression of archaeal ammonia oxidation machinery across disparate environmental and culture conditions

Paul Carini[1,3], Christopher L. Dupont[2], Alyson E. Santoro[1,4]

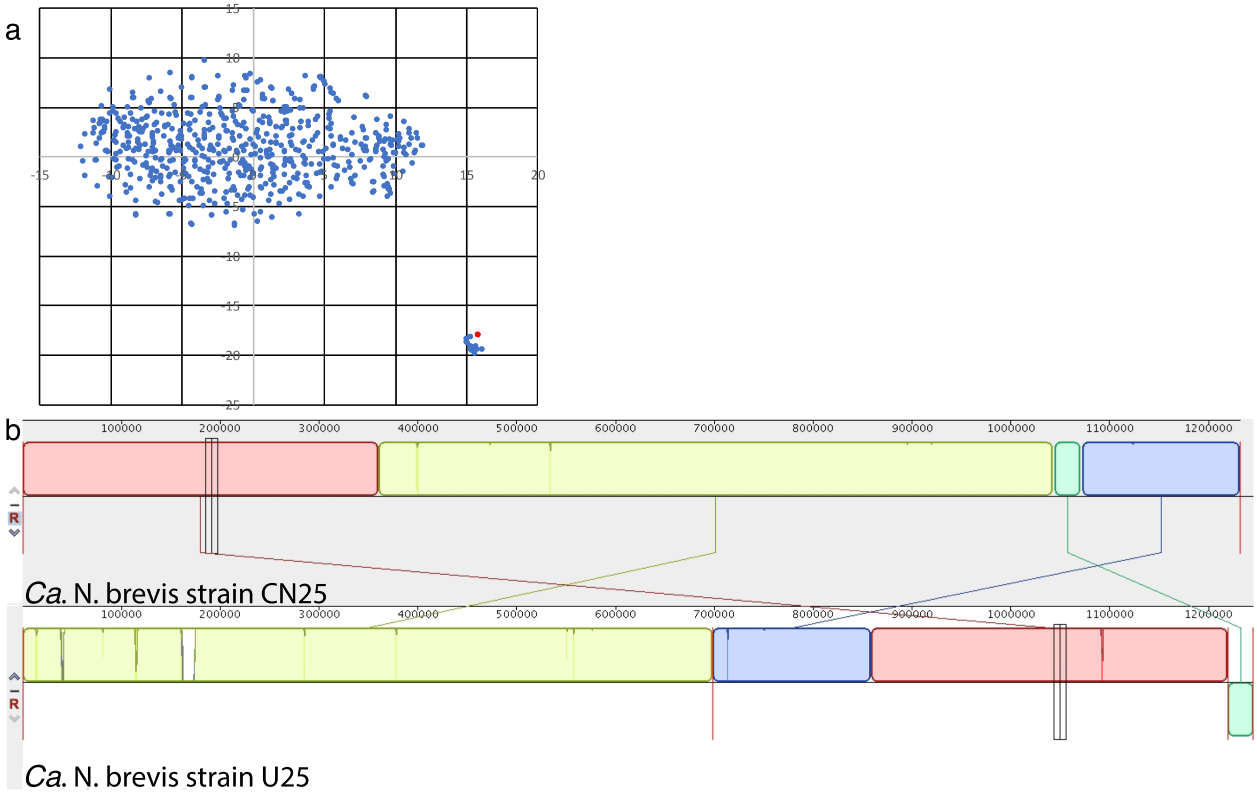


**Supplementary Figure 1:** *Ca*. N. brevis str. U25 metagenome binning and alignment to strain *Ca*. N. brevis str. CN25. (a) Vizbin analysis of 5 mer utilization of the assembly from an thaumarchaea enrichment culture maintained on urea. Contigs containing any thaumarchaea predicted genes are in red and code for the *Ca*. N. brevis str. U25 genome. (b) Mauve alignment of *Ca*. N. brevis strains CN25 (top) and U25 (bottom). The genomes are syntenic with only a few indels between the two strains (listed in Supplementary Table 1) (illustrated as white vertical bands). Red vertical lines in the U25 genome in (b) are contig breaks.

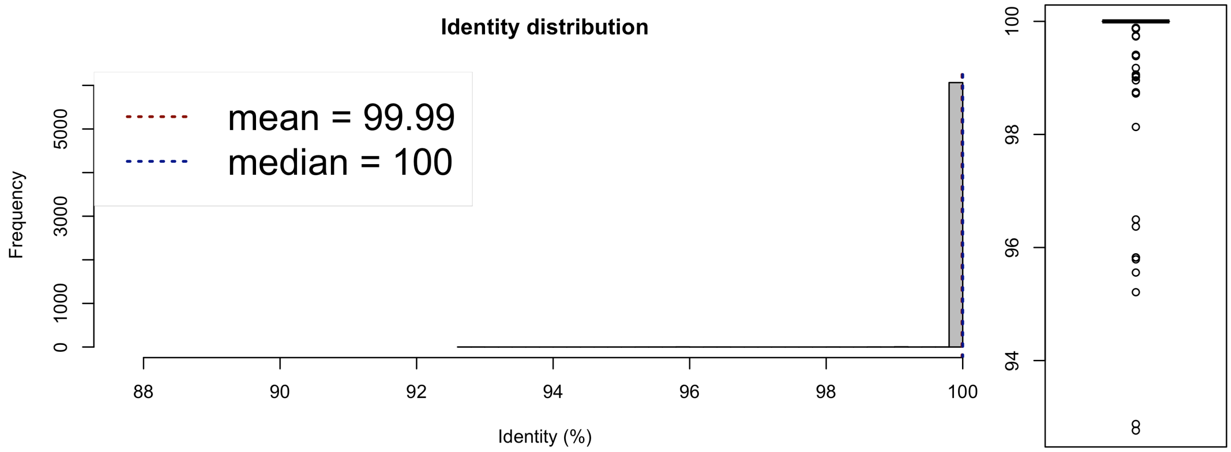


**Supplementary Figure 2:** Simulated DNA-DNA hybridization analysis for the CN25 and U25 genomes using the methods of ref(Goris *et al.*, 2007). The mean nucleotide identity of 99.99% is for a two-way average nucleotide identity analysis. The most divergent conserved genomic segment is 92% nucleotide identity.

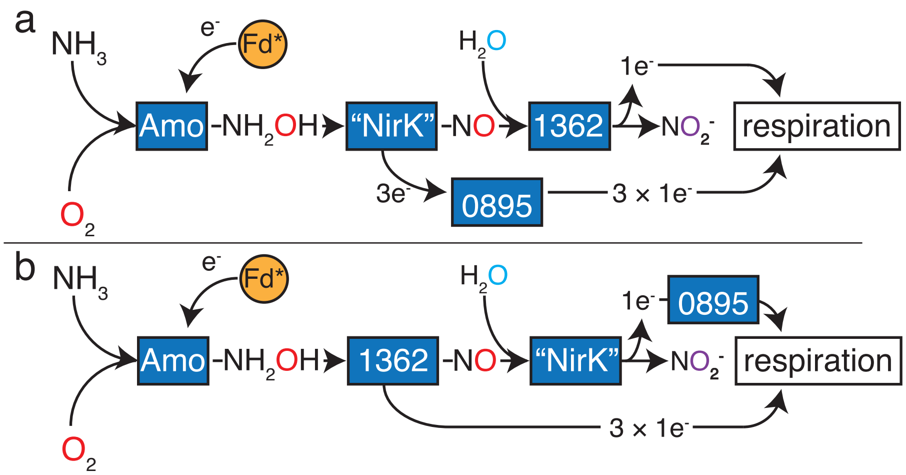


**Supplementary Figure 3:** Proposed ammonia ($NH_3$) oxidation pathways for thaumarchaea based on co-expression of pseudoperiplasmic-localized Cu metalloenzymes. Amo, T478_1362, Fd1, Fd2, "NirK" and T478_0895 are co-expressed in a conserved module (the AMO module) across different environments as illustrated in Fig. 3 in the main text. In these models, a membrane-anchored PEFG-CTERM domain-containing Cu metalloenzyme (T478_1362) acts in concert with "NirK" and T478_0895 to catalyze a two-step reaction: the 3-electron oxidation of hydroxylamine ($NH_2OH$) to nitric oxide (NO) followed by the one-electron oxidation of NO to $NO_2^-$. In (a) "NirK" acts as the hydroxylamine oxidase. The model proposed in (a) is supported by sequence structure threading of T478_1362, which predicts the cupredoxin domain has an open configuration, as in a recently characterized purple cupredoxin from *N. maritimus* that oxidizes NO to $NO_2^-$(Hosseinzadeh *et al.*, 2016). Additionally, the model in (a) is consistent with the potential for NirK to carry three electrons by way of three Cu atoms(Walker *et al.*, 2010); and the potential for $NH_2OH$ to be oxidized to NO by an uncharacterized nitrite reductase (Ritchie and Nicholas, 1972; 1974). In the model depicted in (b), T478_1362 acts as the hydroxylamine oxidase. Model (b) is consistent with the ability of purified bacterial NirK to favor the formation of $NO_2^-$ from NO and water at biological pH(Wijma *et al.*, 2004). Moreover, NirK may catalyze the final step in a three-step ammonia oxidation pathway in ammonia-oxidizing bacteria (AOB)(Caranto and Lancaster, 2017). In both model (a) and (b), electrons are putatively shuttled from NirK to respiratory complexes by membrane-anchored cupredoxin-containing T478_0895. Two ferredoxins (Fd* = Fd1 and Fd2) were also co-expressed with the core ammonia oxidation machinery. These ferredoxins are predicted to be cytoplasmic and may play a role in supplying the electrons required for the initial oxidation of $NH_3$ to $NH_2OH$ by Amo (see main text). Enzyme complexes colored blue are Cu metalloenzymes. Enzymes colored orange are iron-containing. Color of the oxygen atom depicts the source: red = molecular oxygen; Cyan=water; purple=one atom from oxygen and one atom from water. Numbers are locus tags with ‘T478_’ omitted.

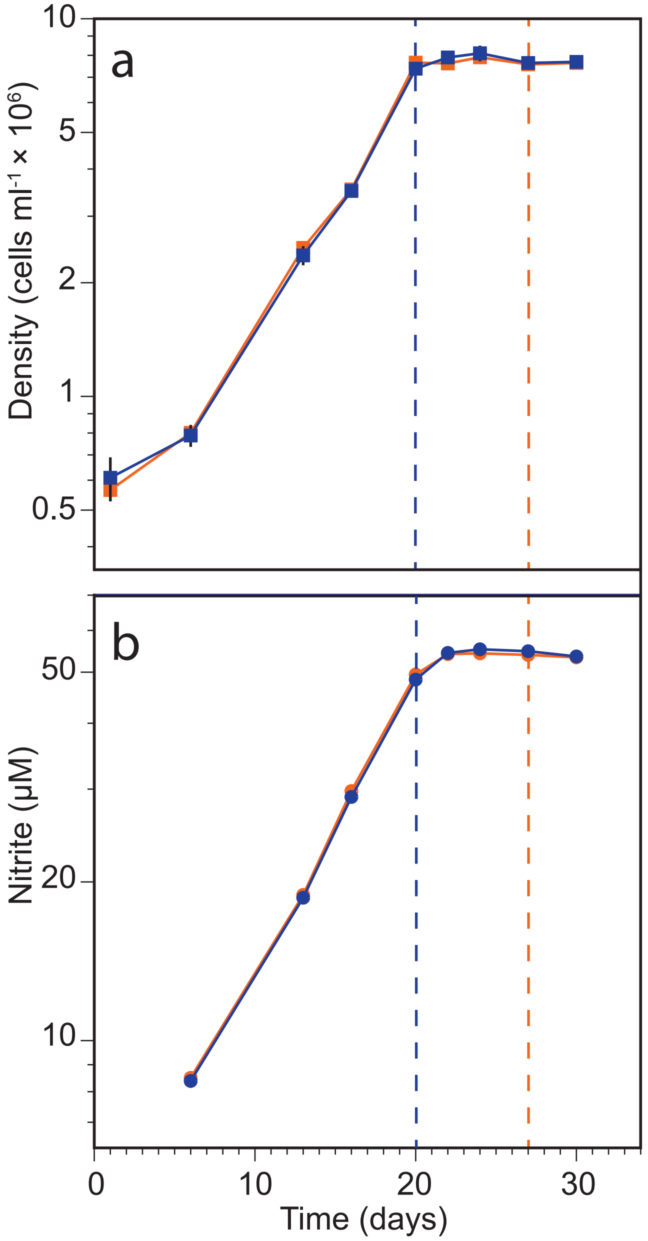


**Supplementary Fig. 4**: *Ca*. N. brevis CN25 growth curves illustrating the average cell density (a) and $NO_2^-$ concentration (b) for cultures harvested for transcriptomes used to investigate the effects of ammonia limitation. Dashed vertical lines are sample time points for exponential phase (blue line) and ammonium-limited stationary phase (orange line). Points are the mean of biological triplicates ± SD. When error bars are not visible, they are smaller than the size of the symbols. Cells were grown in ONP medium(Santoro and Casciotti, 2011) with 50 µM $NH_4Cl$, as described in materials and methods.

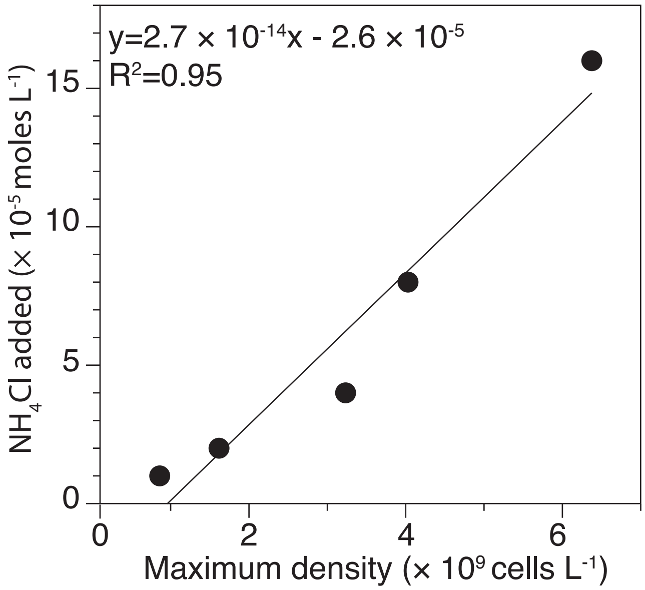


**Supplementary Fig. 5**: *Ca*. N. brevis CN25 molar growth yield in response to ammonium ($NH_4^+$) additions, illustrating that $NH_4^+$ was limiting growth in the transcriptome experiments. Points are the maximum density achieved by *Ca*. N. brevis str. CN25 as a function of $NH_4Cl$ titrations. Linear regression through all five points is shown, with equation and $R^2$. Cells were grown in ONP medium(Santoro and Casciotti, 2011) as described in materials and methods.

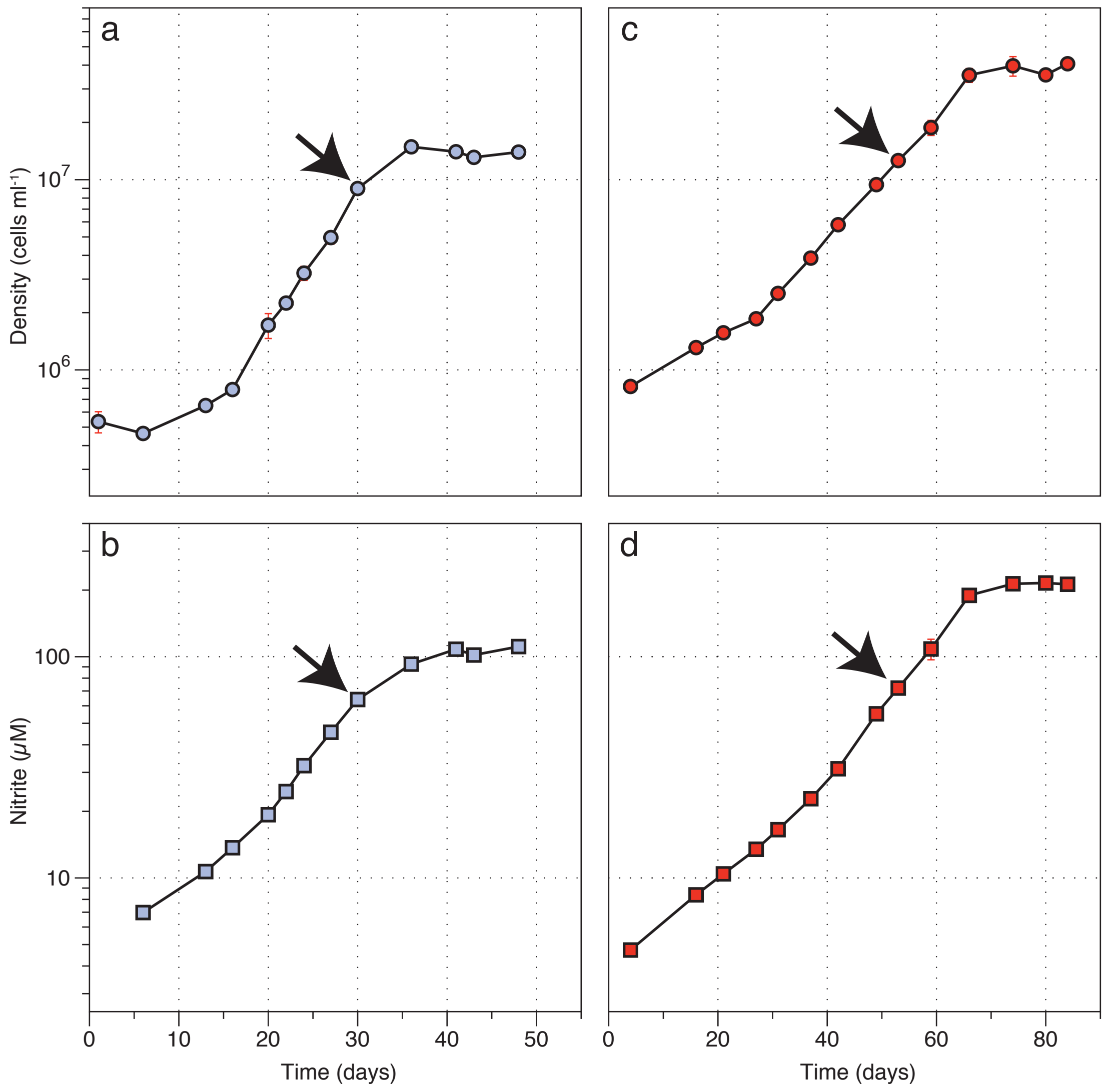


**Supplementary Fig. 6**: *Ca*. N. brevis CN25 (a,b) and U25 (c,d) growth curves illustrating the average cell density (a,c) and $NO_2^-$ concentration (b,d) for cultures harvested for transcriptomes. Arrowheads indicate the cell density and nitrite concentration at the time of sampling. Points are the mean of biological triplicates ± SD. When error bars are not visible, they are smaller than the size of the symbols. Cells were grown in ONP medium (Santoro and Casciotti, 2011) with 100 µM $NH_4Cl$ (a,b) or 100 µM urea (200 µM urea-N) (c,d), as described in materials and methods.

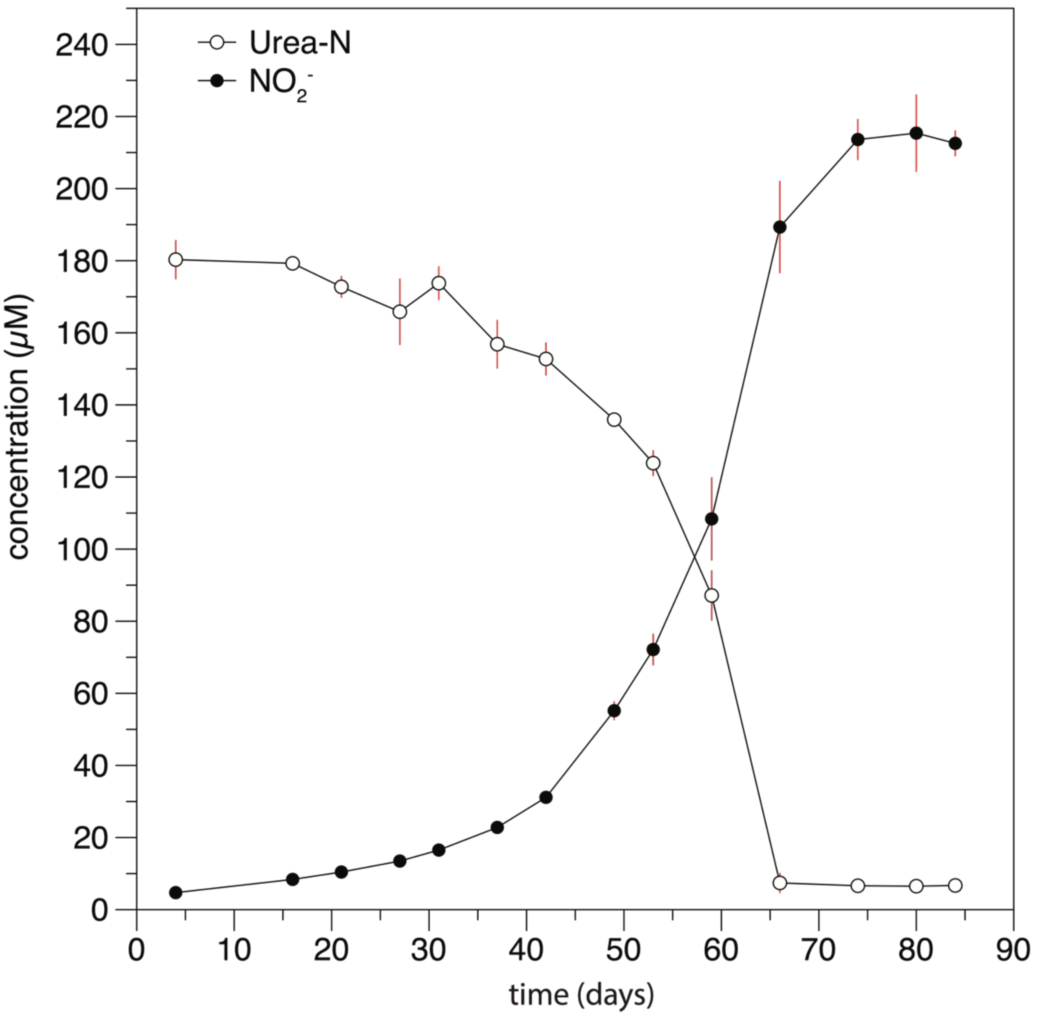


**Supplementary Figure 7:** *Ca*. N. brevis strain U25 oxidizes urea-N to $NO_2^-$. Points are the average ± SD (error bars) concentration of urea-N (open circles) and $NO_2^-$ (filled circles) of triplicate *Ca*. N. brevis str. U25 cultures. Cells were grown in ONP medium(Santoro and Casciotti, 2011) with 100 µM urea, as described in materials and methods.
